# Grey Pineapple Mealybug, *Dysmicoccus neobrevipes* Beardsley: A New Devastating Pest of Tuberose in India

**DOI:** 10.1101/2024.09.24.614774

**Authors:** D. M. Firake, Sunil Joshi, P. Naveen Kumar, T. N. Saha, V.S. Raju Dantuluri, K. V. Prasad

## Abstract

Tuberose (*Agave amica*) is a highly valued flower crop in India, known for its consistently high returns and ability to thrive in various climatic conditions. Severe outbreaks of the grey pineapple mealybug, *Dysmicoccus neobrevipes* Beardsley (Hemiptera: Pseudococcidae), have been observed in several villages of the Pune district, Maharashtra (India), since 2021, leading to significant economic losses. This report provides an illustrative morphological diagnosis of *D. neobrevipes* and basic information on its field establishment, impact on tuberose crops, nature of the damage and field symptoms etc. Surveys conducted in major tuberose-growing areas of Pune district (Maharashtra) revealed that the mealybug primarily infests the underground and basal portion of tuberose plants, causing stunted growth, drooping, and ultimately death of the plants. *D. neobrevipes* produces honeydew, fostering sooty mold growth that impairs photosynthesis, reduces flower yield. It also infests tuberose bulbs, spoiling them in storage. Based on primary scientific literature, this study represents the first scientifically confirmed record of *D. neobrevipes* as a new pest of tuberose in India. The ant species *Solenopsis geminata* was found attending to mealybug colonies, aiding their spread and creating a nuisance for farmers during routine field activities. A total of 87.09% of the surveyed fields (n=62) showed mealybug infestation, ranging from 68% to 97%. Raising awareness among the farmers and implementing regular monitoring in tuberose-growing areas are crucial steps for developing effective management practices and preventing the further spread of this pest to other regions.

---

Tuberose *Agave amica* (Medikus) Thiede & Govaerts (Syn: *Polianthus tuberosa*; Asparagaceae) is widely cultivated across various Indian states for its loose flowers and floral spikes. Tuberose, popularly known as ‘*Rajanigandha* (The Fragrance of the Night) or *Sungandharaja* (King of Fragrance) is considered as the most popular flower crop by small and marginal farmers as it provides consistent and high returns throughout the year^1^. Moreover, it is a hardy crop, capable of withstanding various biotic stresses, and adaptable to a wide range of soils and climatic conditions^2^. Severe outbreaks of this mealybug species were observed in 2021 in the tuberose fields of ICAR-DFR, Pune, as well as in Yavat and the surrounding areas of Pune district, which is a major tuberose-growing region in Maharashtra. Mealybug colonies were found developing on underground parts and basal region of tuberose plants. The affected plants were observed to be completely dried out and drooping, with the infestation being so severe (up to 100%) that farmers were compelled to destroy their entire fields. This mealybug species, including its field characteristics, appeared to differ from the striped mealybug, *Ferrisia virgata* Cockerell (Hemiptera: Pseudococcidae), which is known to damage tuberose crops in India^3,4^. To confirm its identity, mealybug specimens were preserved in 70% ethanol and slide-mounted in Canada balsam, following the procedure described by Watson & Chandler^5^. The terminology for the adult female’s morphological structures is based on the definitions provided by the Williams and Granara de Willink^6^. The morphology of the slide-mounted adult female was examined using 30 specimens mounted on 10 slides at the Division of Germplasm Characterization, ICAR-National Bureau of Agricultural Insect Resources (ICAR-NBAIR), Bengaluru, India. Morphological measurements were recorded using an eyepiece reticle on an Olympus BX 51 microscope, calibrated with a stage micrometer. These measurements represent maximum dimensions, with setal lengths including the base. Slide-mounted adult females were examined using a Nikon Digital Sight DSVI-1 on a Nikon Eclipse 80i microscope, with measurements obtained from digital images using the M205 A Leica Application Suite.

The mealybug species infesting tuberose plants was found to be grey pineapple mealybug, *Dysmicoccus neobrevipes* Beardsley (Hemiptera: Pseudococcidae). Diagnostic characters of *D. neobrevipes* are depicted in Fig.

1. The morphological characters indicated in Fig. 1. are adapted from Williams and Granara de Willink^6^ and Beardsley^7^. *D. neobrevipes* is taxonomically close to the pink pineapple mealybug, *D. brevipes*^6^, and several morphological traits of both species closely resemble each other. Recently, Choudhary et al.^7^ described the morphological characteristics of *D. brevipes* specimens collected from garden pea fields, which show a significant similarity to some of the characters observed in *D. neobrevipes* females collected from tuberose (Fig. 1a to Fig. 1m). These characters include the shape of the adult female’s body, which is oval to broadly oval in shape, with 17 pairs of cerarii (Fig. 1a). The ventral surface of each lobe with a quadrate sclerotized area (Fig. 1b). Each antenna typically consists of 8 segments (Fig. 1c). The legs are well developed, with a stout claw that lacks a denticle (Fig. 1d). Translucent pores are absent from the hind coxa (Fig. 1e), but are abundant on the posterior surfaces of the hind femur (Fig. 1f) and hind tibia (Fig. 1g). The circulus is irregular or shaped like a coffee bean, divided by an intersegmental line (Fig. 1h). The ostioles are well developed, featuring sclerotized inner lips along with small, stout setae and trilocular pores around the periphery (Fig. 1i). Each anal lobe cerarius contains 2 enlarged conical setae and 1 or 2 auxiliary setae (Fig. 1j), along with a compact group of trilocular pores, all located on a more or less circular sclerotized area. The penultimate cerarii (Fig. 1k) are slightly smaller than those on the anal lobes, each comprising 2-4 conical setae, 3-7 auxiliary setae, and a compact group of trilocular pores. The cerarii on the head (Fig. 1l) each have 3-6 conical setae. The anal ring is having 6 setae (Fig. 1m). Other characteristics of female *D. neobrevipes* collected from tuberose are detailed in Fig. 1n to Fig. 1v. The dorsal surface is scattered with short, stiff setae throughout the surface (Fig. 1n). Similar short stiff setae present on surface anterolateral to anal ring (Fig. 1o). Large discoidal pores, each larger than a trilocular pore, present conspicuous medially on the dorsum of the abdomen, particularly on segments VI-VIII (Fig. 1p), each with reticulated surface. Smaller discoidal pores with plain surface, always 1-3 present adjacent to each eye (Fig. 1q). Ventral surface with long flagellate setae around vulva (Fig. 1r) and normal shorter flagellate setae present throughout venter. Multilocular disc pores (Fig. 1s) present around the vulva, at the anterior edge of abdominal segment VII, and the posterior edge of segment VI, in the median areas only. Trilocular pores (Fig. 1t) are evenly distributed, being less numerous on the venter than on the dorsum. Oral collar tubular ducts come in two sizes: a small type (Fig. 1u) is sparsely distributed across the abdominal segments and median area of the thorax, while a larger type (Fig. 1v) is found in groups along the margins of the posterior abdominal segments.

**Fig. 1.**
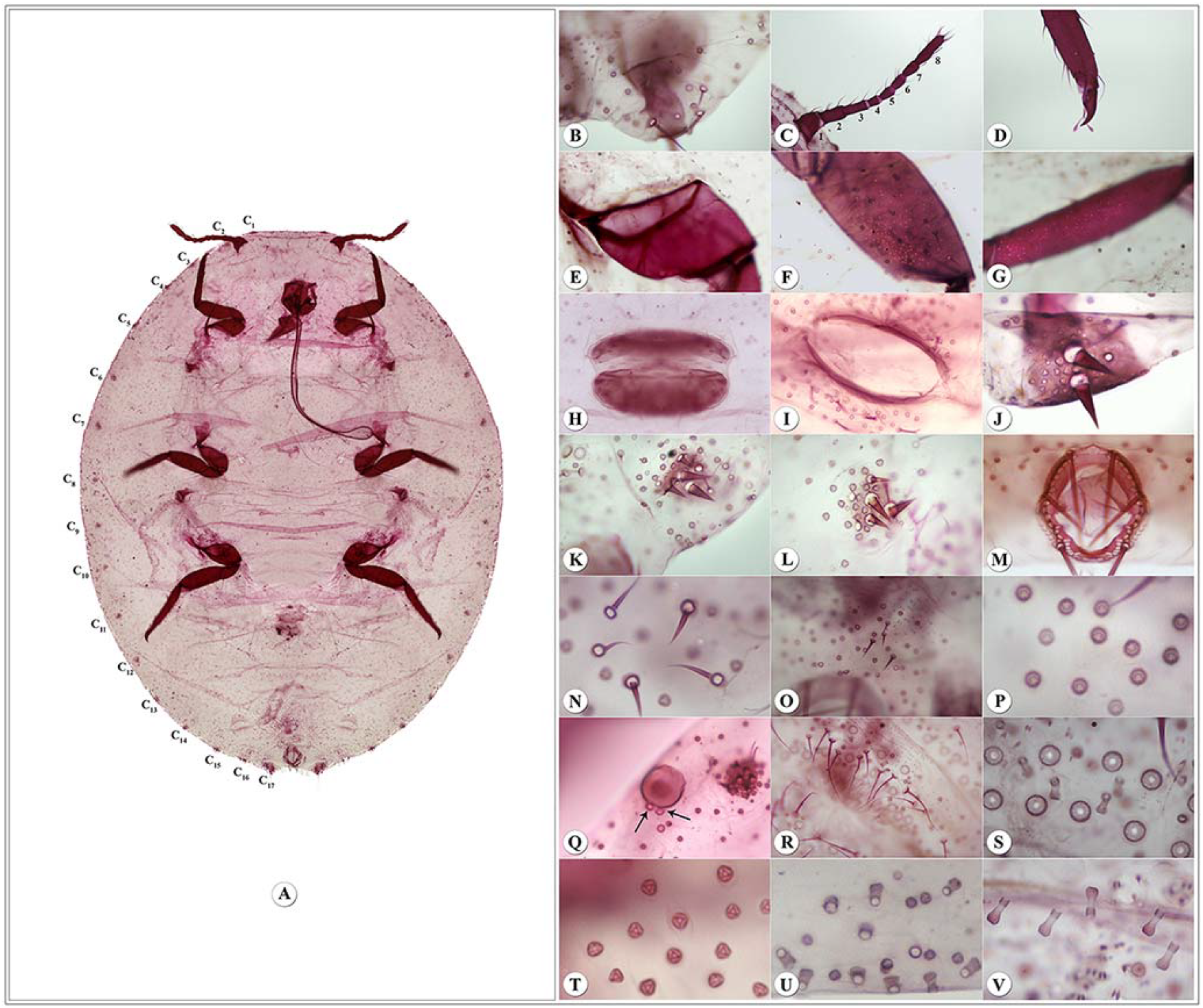
Taxonomic characters of slide mounted female of *Dysmicoccus neobrevipes* Beardsley. Body derm showing 17 cerarii marked as C1 – C17 (a); Ventral surface of anal lobe (b); Antenna (c); Claw (d); Coxa without translucent pores (e); Femur with translucent pores (f); Tibia with translucent pores (g); Circulus (h); Ostiole (i); Anal lobe cerarius (j); Penultimate cerarius (k); Cerarius on head (l); Anal ring (m); Dorsal setae (n); Dorsal setae on abdominal segment VIII anterior to anal ring (0); Dorsal discoidal pores (p); Discoidal pores near eye (q); Ventral setae around vulva (r); Multilocular pores (s); Trilocular pores (t); Smaller oral collar tubular ducts (u); Larger tubular ducts (v)

Originating from pantropical regions, *D. neobrevipes* has now spread to several countries worldwide, including India. This species was first described by Dr. J. W. Beardsley in 1959 from mealybug specimens collected on tuberose (*Polianthes tuberosa*) in Hawaii, USA^8^. *D. neobrevipes* is a polyphagous species associated with plant species across 67 genera in 40 plant families^9^. Although various main host plants of this species have been documented in different countries, pineapple, banana^10^, and sisal (*Agave sisalana*)^11^ are recognized as major hosts. In these crops, *D. neobrevipes* has established itself in the fields and caused significant economic damage. *D. neobrevipes* has been reported in many countries across the Americas, Africa, Europe, Asia, and Oceania. This species is frequently intercepted on various hosts at U.S. ports of entry, including tuberose from Taiwan, and originating from different countries^12^. In Asia, severe outbreaks of *D. neobrevipes* have been observed on various crops in Malaysia, the Philippines, Thailand^13-15^ and China^11^. In India, *D. neobrevipes* has been documented on various crops, including banana^14^. Although this species was originally described from specimens collected from tuberose in the USA, and tuberose is considered as its main host in some countries, there is still a lack of basic information including its natural history on tuberose plants. Given the destructive nature of *D. neobrevipes*, it is imperative to obtain this fundamental information. To address this need, we conducted a detailed investigation and present here an illustrative morphological diagnosis of *D. neobrevipes*, along with essential information on its field establishment, impact on tuberose crops, nature of the damage, and observed field symptoms.

Extensive surveys were conducted in major tuberose-growing localities of Maharashtra during August 2022. Observations on mealybug incidence, damage symptoms, and severity (%) were recorded by counting the number of damaged plants in relation to the total number of plants in 100 sqm area in each field. A total of 62 tuberose fields in Yavat and 10 adjoining villages were examined for mealybug (*D. neobrevipes*) infestation. Among them, 54 (87.09%) tuberose fields were found to have mealybug incidence, with infestation levels ranging from 68% to 100%. Because *D. neobrevipes* hides underground and remains concealed during the early stages of infestation, farmers struggled to control it effectively, even with the use of chemical pesticides, which were often non-systemic. Consequently, many farmers with infested fields were prepared to destroy their entire crops to eliminate the mealybugs.

Numerous mealybug nymphs were observed feeding primarily on the sap of underground parts, leaves and shoots of basal region, which resulted in stunted plant growth. Severely infested plants showed symptoms of drooping, complete blackening of leaves and shoots, and eventually died (Fig.2a to 2c). *D. neobrevipes* impairs the plant’s photosynthetic capacity by excreting sugary honeydew, which fouls plant surface and fosters the growth of sooty mold. This mold blocks sunlight and air from reaching the leaves, further hindering photosynthesis. Additionally, mealybug colonies were found on the spikes of infested plants, leading to poor-quality flowers and reduced yield. Damaged plants often appeared drooped, with rotting underground portions. The infested tuberose fields very often need to be destroyed, as *D. neobrevipes* attacks newly emerging shoots, starting from the underground parts and progressing upward until the shoots die (Fig. 3a). Moreover, *D. neobrevipes* can spread from the field to storage through infested bulbs during harvesting, where they continue to develop and damage the stored tuberose bulbs, making them unsuitable for planting (Fig. 3b & 3c). In infested tuberose fields, the large numbers of fire ants (*Solenopsis geminata*) were observed attending mealybug colonies at nearly all surveyed locations including in stored bulbs. Rarely, but *Camponotus compressus* was also recorded attending colonies of *D. neobrevipes* in few locations. These ants not only protect *D. neobrevipes* from natural enemies but also facilitate its spread by transporting the mealybugs to new plants^16^.

**Fig. 2.**
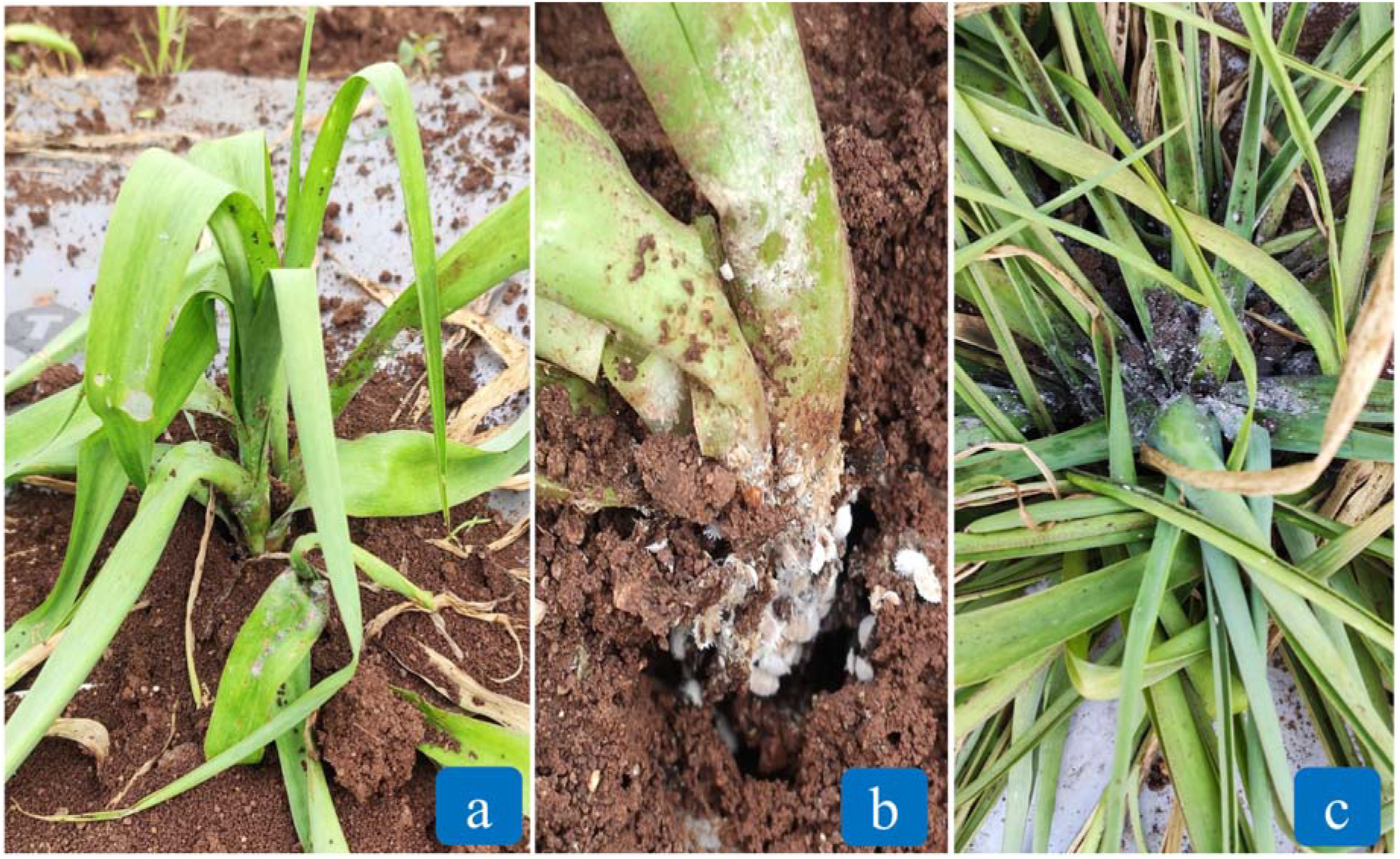
Symptoms of mealybug, *D. neobrevipes* infestation to tuberose plants. Damage symptoms at initial phase of *D. neobrevipes* infestation include a loss of vigor, downward drooping of leaves, and drying of leaf tips (a); *D. neobrevipes* colonies developed on underground parts of the tuberose plants along with large number of ants, *Solenopsis geminata* attending mealybugs (b); *D. neobrevipes* infestation started developing on basal portion and above ground plant parts and excessive honeydew excretion by *D. neobrevipes* leading to blackening of leaves and shoots (c)

**Fig. 3.**
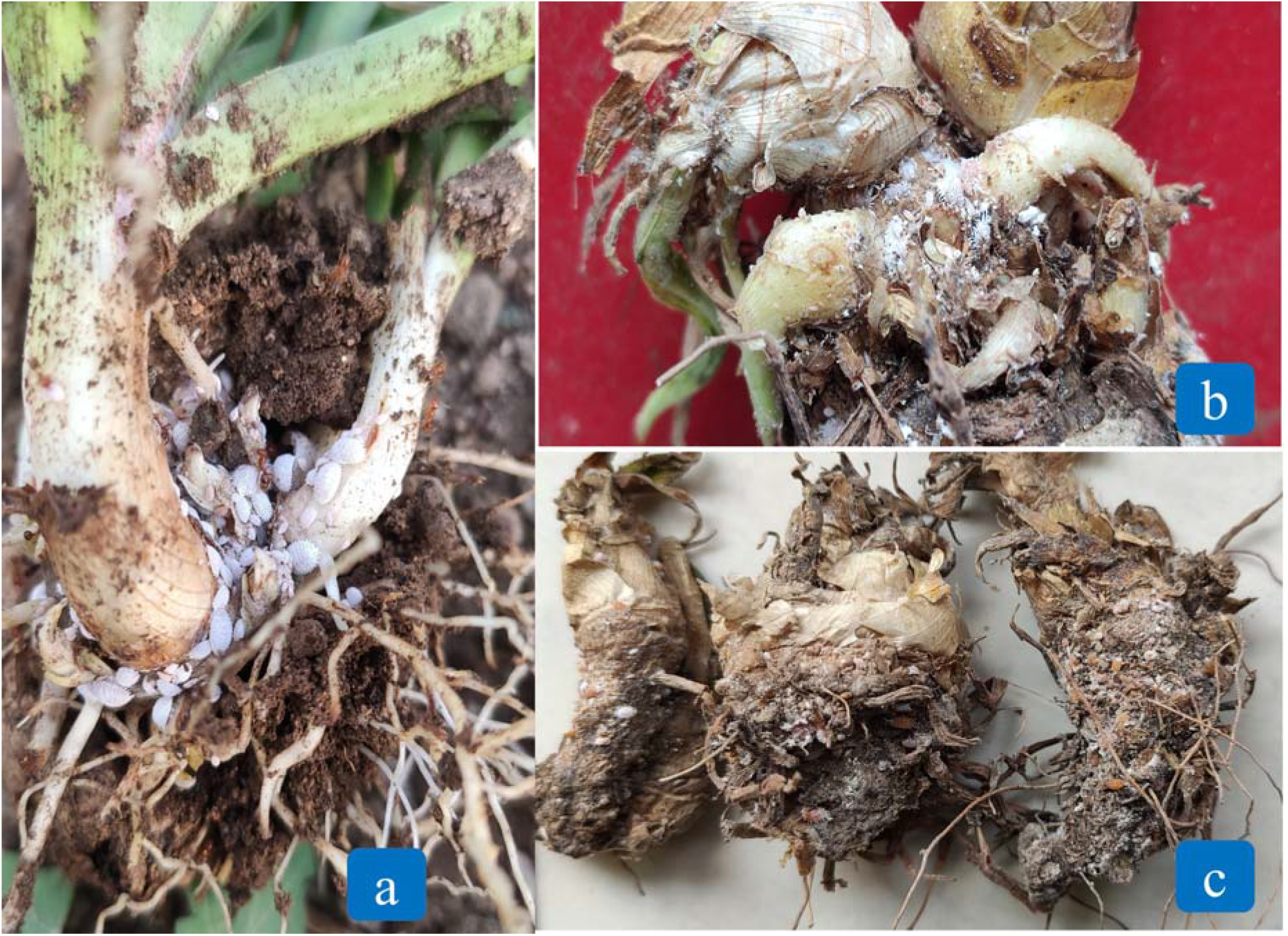
Colonies of mealybug (*D. neobrevipes*) forming on newly emerging shoots, beginning at the underground parts and spreading upward to the shoots and leaves (a); *D. neobrevipes* colonies developing on freshly harvested bulbs (b); and completely damaged buds unsuitable for planting (c).

*D. neobrevipes* reproduces sexually^13^ and is known to cause wilt disease in pineapple in Hawaii. It is closely related to the mealybug *D. brevipes*, though these two species can be differentiated by specific taxonomic features. Besides morphological differences, *D. neobrevipes* primarily targets the aerial parts of plants, while *D. brevipes* feeds on the roots and basal portions of host plant^8^. However, we observed that *D. neobrevipes* infestations initially start in the underground portions of tuberose plants, with mealybug colonies predominantly found on the basal parts. These colonies can also be found on the aerial parts of the plant, including the shoots, leaves, and even spikes during flowering and post flowering stage. In the case of pineapple, *D. neobrevipes* is known to primarily attack the aerial parts of the plants, unlike its close relative *D. brevipes*, which mostly develops in the underground portions^8^. Even when infesting aerial parts, *D. neobrevipes* tends to conceal itself in protected areas of the pineapple, such as under bracts, making visual detection challenging^18^. This behavior suggests that *D. neobrevipes* prefers to inhabit concealed areas of plant structures, which may not be available before spike emergence in tuberose. Consequently, the initial infestation in tuberose were primarily found in the underground and basal portions of the plant. Based on primary scientific literature, this study represents the first scientifically confirmed report of *D. neobrevipes* as a significant pest of tuberose.

The present study indicates that *D. neobrevipes* has emerged as a new devastating pest of tuberose in Maharashtra, India. Its concealed habit during the initial stages of infestation makes early visual detection challenging. Since ants facilitate the spread of this pest and its protection from natural enemies, managing ant populations should be a primary focus in controlling mealybugs. Additionally, *D. neobrevipes* can spread with infested bulbs and they continue to develop in storage, rendering the bulbs unsuitable for planting. Chances of spread of mealybugs through infested planting material will also be very high. Therefore, it is recommended to thoroughly wash harvested bulbs with a 1% detergent solution and allow them to dry in the shade before storage. Stored bulbs should be regularly inspected for mealybug colonies and washed again with 1% detergent before planting to prevent initial field infestations. To tackle the issue of this emerging pest affecting tuberose, it is essential to focus on increasing awareness among farmers and conducting regular monitoring in tuberose-growing regions throughout India. Further efforts are needed to understand its field bio-ecology and natural enemies to develop effective management practices

## Acknowledgements

The funding for the institute project “Eco-friendly Pest Management in Commercial Loose Flower Crops, Project code: 45/S/IPP/12” by the Indian Council of Agricultural Research, New Delhi is duly acknowledged. Authors sincerely acknowledge Dr. Himender Bharati, Professor, Punjabi University, Patiala (India) for identification of ant specimens associated with *D. neobrevipes*

## Statements and Declarations

Authors declare that this work has not been published previously or it is not under consideration for publication elsewhere. The article’s publication is approved by all the authors and tacitly or explicitly by the responsible authorities where the work was carried out

## Conflicts of interest statement

The authors declare that they have no known competing financial interests or personal relationships that could have appeared to influence the work reported in this paper.

## Notes

### Competing Interest Statement

The authors have declared no competing interest.

